# Mother centrioles are dispensable for deuterosome formation and function during basal body amplification

**DOI:** 10.1101/373662

**Authors:** Huijie Zhao, Qingxia Chen, Qiongping Huang, Xiumin Yan, Xueliang Zhu

**Affiliations:** State Key Laboratory of Cell Biology, CAS Center for Excellence in Molecular Cell Science, Shanghai Institute of Biochemistry and Cell Biology, Chinese Academy of Sciences; University of Chinese Academy of Sciences, 320 Yueyang Road, Shanghai 200031, China; School of Life Science and Technology, ShanghaiTech University, 100 Haike Road, Shanghai 201210, China

**Keywords:** basal body, centriole, deuterosome, multicilia, Plk4

## Abstract

Mammalian epithelial cells use a pair of mother centrioles (MCs) and numerous deuterosomes as platforms for efficient basal body assembly during multiciliogenesis. How deuterosomes form and function, however, remain controversial. They are proposed to either arise spontaneously followed by maturation into larger ones with increased procentriole-producing capacity or be assembled solely on the young MC, nucleate procentrioles under the MC’s guidance, and released as procentriole-occupied "halos". Here we show that both MCs are dispensable for deuterosome formation in multiciliate cells. In both mouse tracheal epithelial and ependymal cells (mTECs and mEPCs), discrete deuterosomes in the cytoplasm were initially procentriole-free and then grew into halos. More importantly, eliminating the young MC or both MCs in proliferating precursor cells through shRNA-mediated depletion of Plk4, a kinase essential to procentriole assembly, did not abolish deuterosome formation when these cells were induced to differentiate into mEPCs. The average deuterosome numbers per cell only reduced by 21% as compared to control mEPCs. Therefore, MC is not essential to the assembly of both deuterosomes and deuterosome-mediated procentrioles.

## Introduction

Epithelial tissues such as those in mammalian trachea, ependyma, and oviduct are abundant in terminally differentiated cells with dense motile cilia. These multiciliated cells each require up to hundreds of centrioles to serve as basal bodies of their cilia (Brooks & Wallingford, 2014, Ishikawa & Marshall, 2011, Spassky & Meunier, 2017). To achieve this, each of the two MCs produces multiple daughter centrioles, breaking the general tight control on the centriole duplication. Furthermore, dozens of deuterosomes emerge as alternative organelles of centriole biogenesis to generate the majority of the basal bodies (Anderson & Brenner, 1971, Dirksen, 1971, Sorokin, 1968). We have previously found that the MC-mediated centriole amplification still uses the canonical ring-shaped platform around its basolateral wall, which contains the Cep63-Cep152-Plk4 complex and other components (Banterle & Gonczy, 2017, Brown, Marjanovic et al., 2013, Habedanck, Stierhof et al., 2005, Hatch, Kulukian et al., 2010, Nigg & Holland, 2018, Sir, Barr et al., 2011) but is hyperactivated through overexpression of its key regulators (Yan, Zhao et al., 2016, Zhao, Zhu et al., 2013), similar to what have been demonstrated in cycling cells by overexpressing Plk4, Cep152, SAS6, or STIL (Arquint, Sonnen et al., 2012, Dzhindzhev, Yu et al., 2010, Kleylein-Sohn, Westendorf et al., 2007, Strnad, Leidel et al., 2007, Vulprecht, David et al., 2012). The deuterosome, on the other hand, adapts Deup1, a paralog of Cep63, to create a similar but MC-free platform of centriole biogenesis (Tang, 2013, Yan et al., 2016, Zhao et al., 2013). Fully assembled centrioles are eventually released from their "cradles" by APC/C-activated proteolysis and mature into basal bodies (Al Jord, Shihavuddin et al., 2017, Zhao et al., 2013).

Deuterosomes have been thought to form autonomously and mediate MC-independent, or *de novo*, centriole biogenesis (Beisson & Wright, 2003, Dirksen, 1971, Nigg & Stearns, 2011, Sorokin, 1968). Consistently, our results suggest that in mTECs deuterosomes are self-assembled, initially nucleating 1-2 procentrioles, and then grow into larger ones with more procentrioles (Fig. 1a) (Zhao et al., 2013). The MCs nucleate their own daughter centrioles during the deuterosome formation (Fig. 1a) (Zhao et al., 2013). On the other hand, a recent study using mEPCs proposes that both deuterosome formation and deuterosome-mediated procentriole nucleation require the young MC: dueterosomes are sequentially assembled at the basolateral side of the young MC; each growing deuterosome associates with more and more procentrioles nucleated under the guidance of the MC until it is released as a procentriole-occupied "halo" after reaching the full size (Fig. 1b) (Al Jord, Lemaitre et al., 2014, Meunier & Spassky, 2016). The MCs nucleate their own procentrioles only after the last halo is released (Fig. 1b) (Al Jord et al., 2014).

**Figure 1.**
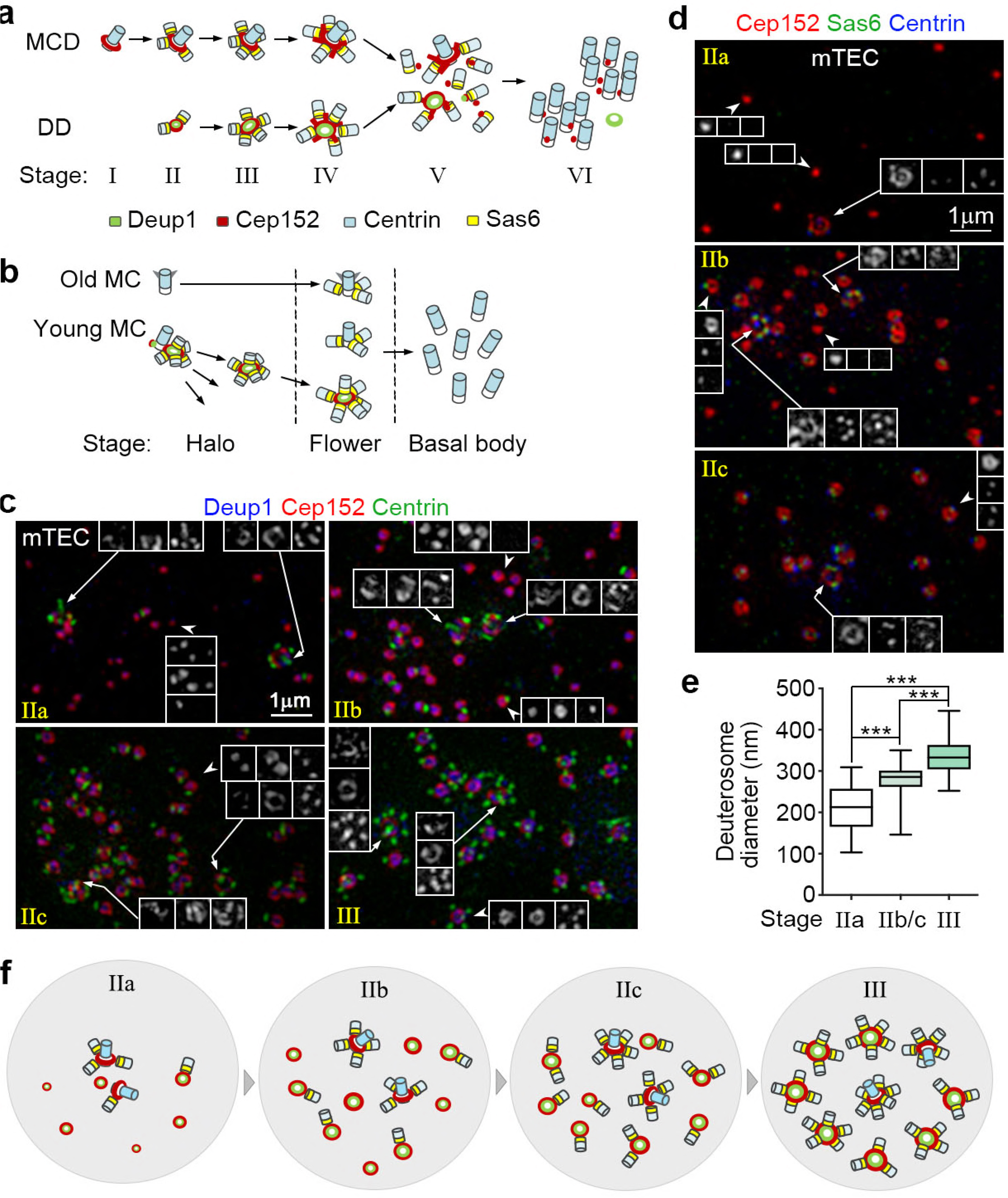
Discrete deuterosomes in mTECs are initially free of procentrioles. **(a,b)** Two current models for the process of massive basal body formation based on studies in mTECs (**a**) and mEPCs (**b**). In (**a**), deuterosomes are proposed to form spontaneously in stage II to each support the assembly of 1-2 procentrioles. They grow in size and nucleate more procentrioles in stage III. Each mother centriole (MC) also nucleates multiple procentrioles from stage II. Stage IV is characterized by the emergence of Cep152-positive protrusions, stage V by the releasing of basal bodies from both deuterosomes and MCs, and stage VI by the formation of basal body clusters under the apical side of the cell membrane (Zhao et al., 2013). In (**b**), deuterosomes are assembled at the lateral wall of the young MC and released one by one as procentriole-occupied “halos” (the halo stage). When the last halo is released, MCs start to generate their own procentrioles. Then the procentrioles on both deuterosomes and MCs start to mature simultaneously (the flower stage) and are eventually released as basal bodies (the basal body stage) (Al Jord et al., 2014). Deup1, Cep152, Centrin, and Sas6 are used as markers respectively for the deuterosome, MC and deuterosome, centriole, and procentriole. **(c,d)** Representative mTECs containing early deuterosomes. mTECs cultured at an air-liquid interface for 2 days (ALI d2) were immunostained for Cep152, Centrin, and Deup1 (**c**) or Sas6 (**d**) and subjected to 3D-SIM. MCs and typical deuterosomes are denoted by arrows and arrowheads, respectively. IIa: MCs contain procentrioles; most deuterosomes are procentriole-free and usually small. IIb: a substantial portion of deuterosomes contains 1 procentriole; deuterosomes are usually medium in size. IIc: most deuterosomes contain 1-2 procentrioles; deuterosomes are usually medium in size. A stage-III cell, which contained larger deuterosomes associated with more procentrioles, is shown in (**c**) for comparison. **(e)** Deuterosome-size distributions in stage IIa, IIb/c, and III. The diameters of at least 348 deuterosomes from 12 mTECs were measured for each group. The bottom and top of the box represent the 25th and 75th percentiles, respectively. The band is the median. The ends of the whiskers indicate the maximum and minimum of the data. Student’s *t* test: ^04-Dec-18***^ *P*<0.001. **(f)** An illustration model for the progression of mTECs from stage IIa to III.

In this study, we examined both mTECs and mEPCs to clarify the discrepancies.

## Results and discussion

### Deuterosomes in mTECs are capable of *de novo* centriole biogenesis

To clarify whether mTECs also adapted the proposed MC-dependent deuterosome production (Fig. 1b), we examined their early deuterosomes. mTECs can be induced to differentiate into multiciliated cells efficiently by culturing at an air-liquid interface (ALI) (You, Richer et al., 2002, Zhao et al., 2013). As mTECs thus cultured for 3 days (ALI d3) were mainly at late stages of the basal body production, we examined those at ALI d2 and found abundant stage-II cells, i.e., cells with deuterosomes that are smaller in size and contain fewer (usually 0-2) procentrioles as compared to those in stage III (Fig. 1c; also refer to the diagrams in Fig. 1a) (Zhao et al., 2013).

Consistent with our previous report (Zhao et al., 2013), both the old and young MCs carried procentrioles in these stage-II mTECs when they were both visible (Fig. 1c,d). By contrast, the size and number of the deuterosomes and the status of their procentrioles, which were better defined by using both Sas6 and Centrin as markers (Fig. 1d) (Leidel, Delattre et al., 2005, Nakazawa, Hiraki et al., 2007), varied dramatically. In a portion of the cells, for instance, deuterosomes were sparse, small (ϕ = 211.1 ± 59.9 nm), and usually procentriole-free (Fig. 1c-e: IIa), suggesting that they are in the early phase of the assembly (Fig. 1f: IIa). In the remaining stage-II cells, deuterosomes were generally larger (ϕ = 277.6 ± 46.2 nm) and more abundant (Fig. 1c-e: IIb/c). Those mingled with deuterosomes without procentriole or with only one procentriole were presumably in the middle of stage II (Fig. 2c-f: IIb). Sometimes deuterosomes of both small and large sizes were observed in these cells (Fig. 1d: IIb). By contrast, those with deuterosomes that were commonly associated with 1-2 procentrioles were in the late stage II (Fig. 1c- f: IIc). In comparison, in stage-III mTECs deuterosomes were 336.1 ± 50.9 nm in diameter and frequently associated with 3-5 procentrioles (Fig. 1c,e, f). These results still support the model of *de novo* centriole biogenesis (Fig. 1a) (Zhao et al., 2013) but not the MC-dependent one (Fig. 1b) (Al Jord et al., 2014).

**Figure 2.**
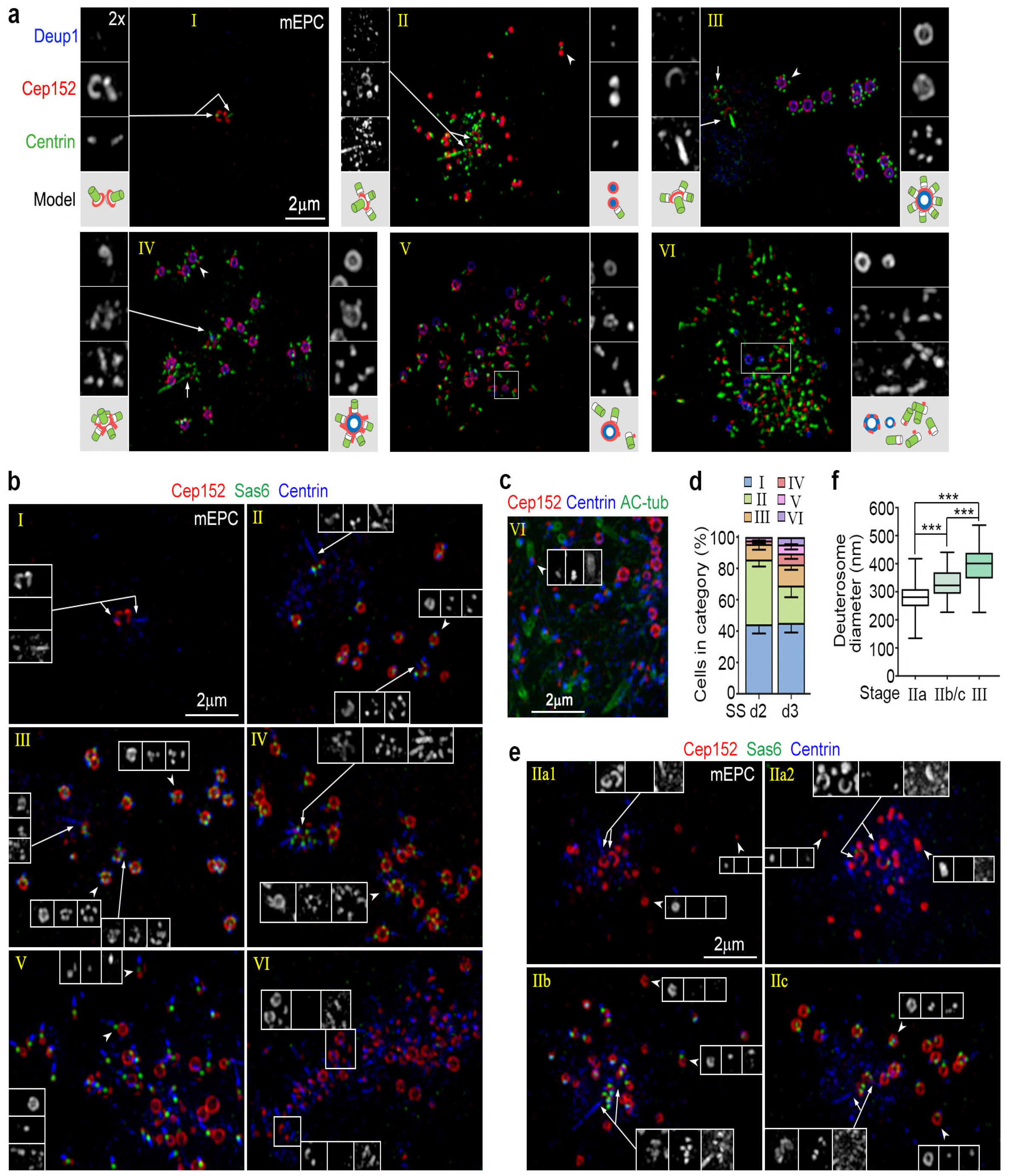
mEPCs resemble mTECs in centriole amplification. **(a)** Typical mEPCs representing different stages of centriole amplification. mEPCs cultured under serum starvation for 3 days (SS d3) were immunostained for Deup1, Cep152, and Centrin, followed by imaging with 3D-SIM. MCs and typical deuterosomes are denoted by arrows and arrowheads, respectively. An illustration is provided for each set of the magnified images. **(b)** Typical mEPCs at SS d3, immunostained for Cep152, Sas6, and Centrin. **(c)** Multicilia formation in a typical stage-VI mEPC. Acetylated tubulin (AC-tub) was used as ciliary marker. Note that the ciliogenesis is asynchronous. **(d)** Stage-distributions of mEPCs at SS d2 and d3. The results were from three independent experiments. At least 108 cells were scored in each experiment and condition. **(e)** Discrete deuterosomes in early stage-II mEPCs were also procentriole-free. mEPCs at SS d2 were immunostained for Cep152, Sas6, and Centrin. **(f)** Deuterosome-size distributions. The diameters of at least 496 deuterosomes from 32 mEPCs were measured for each group. Student’s *t* test: ^***^ *P*<0.001.

### Centriole amplification in mEPCs resembles mTECs

Next we investigated whether deuterosomes in mEPCs are produced differently from those in mTECs. Cultured radial glia isolated from mouse brain tissues can be induced to differentiate into multiciliated mEPCs through serum starvation (Delgehyr, Meunier et al., 2015, Spassky, Merkle et al., 2005). We examined the mEPCs at day 2 or day 3 post serum starvation (SS d2 or d3) because they were undergoing active centriole amplification (Al Jord et al., 2014, Zheng, Liu et al., 2018). As somehow Centrin-positive aggregates were frequently observed in mEPCs, especially in the area around the MCs, we also used Sas6 as additional procentriole marker (Leidel et al., 2005, Nakazawa et al., 2007).

We found that the mEPCs could also be grouped into six stages (Fig. 2a,b), similar to mTECs (see Fig. 1a) (Zhao et al., 2013): (1) those containing only a pair of Cep152-positive, Deup1-negative centrioles were in stage I; (2) those containing small deuterosomes with mostly 0-2 procentrioles were in stage II; (3) those in the "halo" or "flower" stages (Al Jord et al., 2014) contained larger deuterosomes commonly with 3-7 procentrioles and could be assigned to stage III or IV, respectively. The stage-IV cells were still distinguished from those in stage III by the emergence of multiple Cep152-positive protrusions from both deuterosomes and MCs. Their Centrin-staining was also elongated as compared to the punctual staining in the stage-III cells; (4) those with partially released basal bodies from their cradles were in stage V. Their basal bodies still contained the Sas6-positive puncta; and (5) those with fully released basal bodies negative for the Sas6 staining were in stage VI (Fig. 2a,b). Similar to mTECs (Zhao et al., 2013, Zheng et al., 2018), stage-VI mEPCs also underwent multiciliogenesis (Fig. 2c). Only their basal bodies did not group into clusters (Fig. 2a-c) as those do in mTECs (Zhao et al., 2013).

Quantifications indicated that 44.3 ± 3.5% (SS d2) and 45.3±3.9% (SS d3) of the cells were morphologically in stage I (Fig. 2d). In addition to those that would soon undergo deuterosome formation, this population also contained cells of other fates because 30-60% of mEPCs were multiciliated at SS d5 or later (Zheng et al., 2018). The mEPCs were more abundant in those at early stages at SS d2 than at SS d3. For instance, stage-II cells occupied 41.3 ± 2.7% at SS d2 but 23.8 ± 5.0% at SS d3 (Fig. 2d). On the other hand, 17.4% of the mEPCs were in stages IV-VI at SS d3, whereas the value was 4.1% at SS d2 and in fact no stage-VI cells were observed (Fig. 2d). These results further support the conclusion that mEPCs progress from stage I to VI during their differentiation into multiciliated cells.

For further clues on early phases of the deuterosome formation, we examined the stage-II mEPCs in detail. We found that, similar to mTECs (Fig. 1), stage-II mEPCs could also be grouped into those in IIa, IIb, and IIc (Fig. 2e). The average deuterosome diameters were 277.5 ± 61.0 nm (IIa) and 330.5 ± 60.5 nm (IIb/c), which were smaller than those in stage-III mEPCs (391.9 ± 76.3 nm) (Fig. 2f). Furthermore, when both the young and old MCs were visible in the mEPCs at stage IIb or IIc, they all bore procentrioles (Fig. 2e; also see Fig. 2a,b). This was also true for the cells in stage III or IV (Fig. 2a,b). In the stage-IIa mEPCs, however, MCs that were with or without procentrioles were observed (Fig. 2e, IIa1 and IIa2).

### MC is dispensable for deuterosome formation

Next we assessed the proposed role of the young MC to deuterosome formation (Fig. 1b) (Al Jord et al., 2014). We reasoned that, as Plk4 is essential for centriole duplication in both cycling and multiciliated cells (Bettencourt-Dias, Rodrigues-Martins et al., 2005, Habedanck et al., 2005, Kleylein-Sohn et al., 2007, Zhao et al., 2013), its depletion in proliferating murine radial glia, stem cells of multiciliate mEPCs (Delgehyr et al., 2015, Spassky et al., 2005), would result in the cells with either one old MC (1MC) or no MC (0MC) (Fig. 3a). Inducing their differentiation by serum starvation would allow us to examine how deuterosome formation is affected by MCs (Fig. 3a).

**Figure 3.**
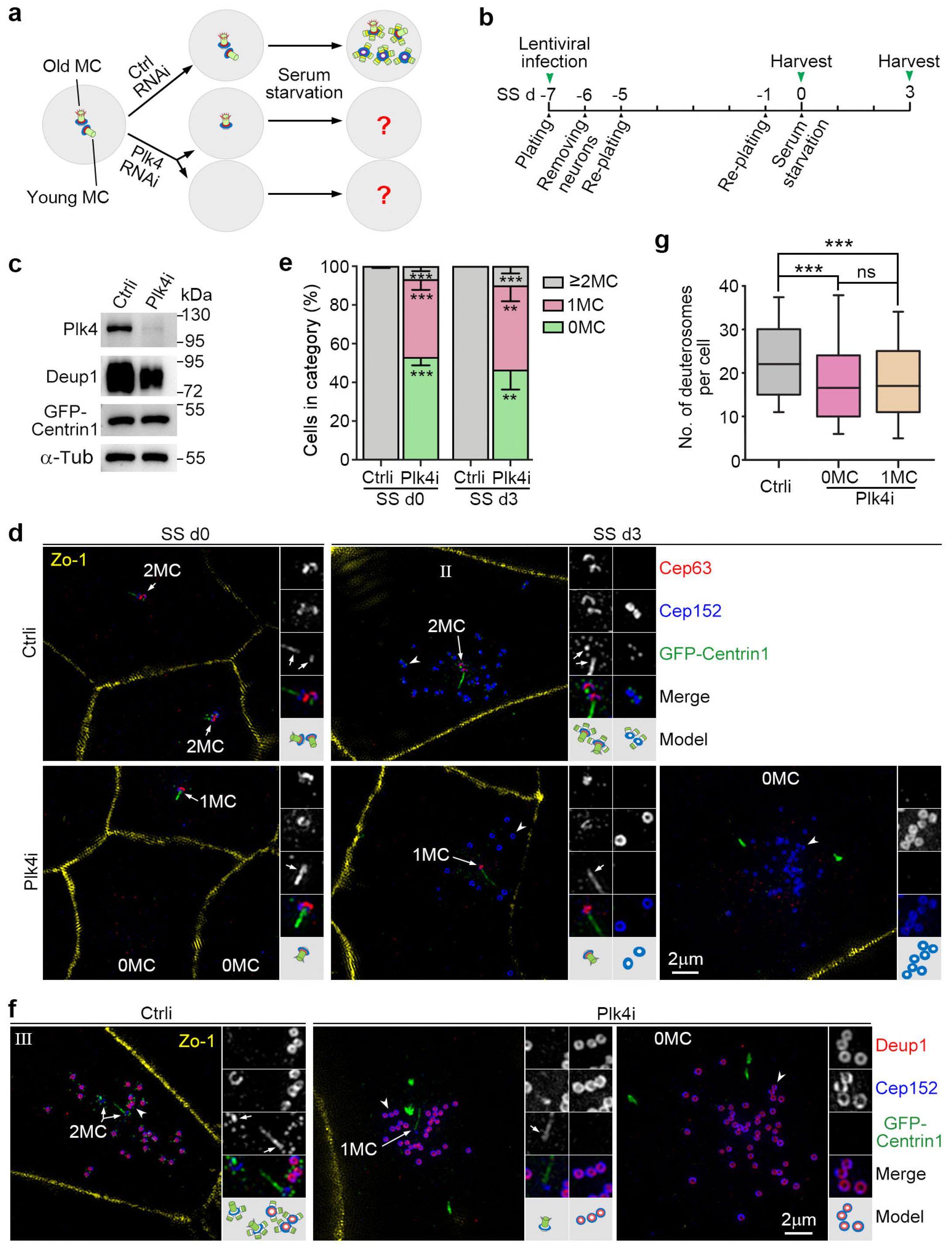
Deuterosomes form efficiently in the absence of MCs. **(a,b)** Experimental design. Depletion of Plk4 is known to abolish centriole duplication in proliferating cells and result in cells with only the old MC (1MC) or no MC (0MC), which could thus be used to examine the contribution of MCs to the deuterosome formation (**a**). We transfected radial glia from mouse brain tissues with lentiviral particles at SS −7 to silence Plk4 expression and examined their progeny cells at SS d0 and d3 (**b**). **(c)** Confirmation of the Plk4 RNAi efficiency using mEPCs at SS d3. The reduced expression of Deup1 in the Plk4-depleted cells is attributed to reduced multiciliate cell differentiation. **(d)** Typical cells at SS d0 and d3, immunostained for Zo-1, Cep152, and Cep63, an MC-specific marker. GFP-Centrin1 expressed from the lentiviruses served as both infection and centriole markers. The green fluorescence was enhanced using anti-GFP antibody and Alexa Fluor-488-conjugated secondary antibody. MCs and representative deuterosomes, denoted respectively by arrows and arrowheads, were magnified 2-fold to show details. An illustration is provided for each set of the magnified images. **(e)** MC contents of the cells examined at SS d0 and d3. The results were from three independent experiments and presented as mean ± SD. At least 186 cells at SS d0 and 80 deuterosome-containing cells at SS d3 were scored in each experiment and condition. Student’s *t* test: ^**^*P*<0.01; ^***^ *P*<0.001. **(f)** Confirmation of deuterosome formation in Plk4i-expressing mEPCs at SS d3 by co-staining for Deup1**. (g)** Box plots for deuterosome numbers per cell. At least 156 deuterosome-containing cells from three independent experiments were scored. Student’s *t* test: ns, not significant; ^***^ *P*<0.001.

We have previously established a lentivirus that can stably co-express an infection marker GFP-Centrin1 with Plk4i, a short hairpin RNA (shRNA) against the murine Plk4 mRNA (Zhao et al., 2013). We found that infecting the glial cells for 7 days before serum starvation (SS d-7) with the virus efficiently depleted Plk4 (Fig. 3b,c). When Zo-1, a tight junction protein (Stevenson, Siliciano et al., 1986), was used to mark cell boundaries, co-staining Cep152 with an MC-specific marker, Cep63 (Brown et al., 2013, Lukinavicius, Lavogina et al., 2013, Zhao et al., 2013), indeed revealed loss of one or both MCs as a common phenotype at SS d0 (Fig. 3d). Quantification indicated an average of 53.0% of 0MC- or 40.2% of 1MC-cells (Fig. 3e). In comparison, 99.8% of the cells infected with a lentivirus co-expressing a control shRNA (Ctrli) with GFP-Centrin1 contained two or more MCs (≥2MC) (Fig. 3e).

Notably, 33.6% of the Plk4i-expressing mEPCs at SS d3 (n=113) were found to still contain deuterosomes (Fig. 3d), which were further confirmed by co-staining with Deup1 (Fig. 3f). As expected, procentriole formation was abolished in these cells (Fig. 3d,f). More importantly, 46.5% of the deuterosome-producing cells had no MC, whereas 43.5% of them contained a single MC (Fig. 3d-f). Only a small portion (10.0%) contained two or more MCs (Fig. 3e). In comparison, deuterosome-containing cells occupied 52.1% of the Ctrli-expressing mEPCs at SS d3 (n=71). Quantifications revealed that deuterosome numbers in the 0MC- and 1MC-mEPCs were very similar (Fig. 3g). The average numbers were 19.3 ± 13.2 (n=172 cells) and 19.2 ± 12.4 (n=156 cells), respectively. Deuterosome numbers increased slightly in the control mEPCs (Fig. 3g), with an average of 23.8 ± 11.5 (n=316 cells). These results indicate that MC is not important for deuterosome formation. The reduced percentage of the deuterosome-containing cells correlated with the reduced protein levels of Deup1 (Fig. 3c), suggesting a decreased differentiation potency of the SS d0-cells, possibly due to Plk4 depletion-induced self-renewal defects of the radial glia (Martin, Ahmad et al., 2014).

To confirm that the 1MC-cells indeed contained only the old MC, we used Cep164 and Odf2, which localize to centriolar appendages specific for the old MC (Graser, Stierhof et al., 2007, Mori, Hazan et al., 2017, Yang, Chong et al., 2018), as markers. In the MC pairs of the Ctrli-expressing cells at SS d0, one was positive for Cep164 and Odf2 at the distal centriolar region, as compared to the proximal localization of Cep152, and the other was negative (Fig. 4a,b). By contrast, the remaining MC in the Plki-expressing cells was always positive for Cep164 and Odf2 at both SS d0 and SS d3 (Fig. 4a,b). Thus, this remaining MC is indeed the old MC.

**Figure 4.**
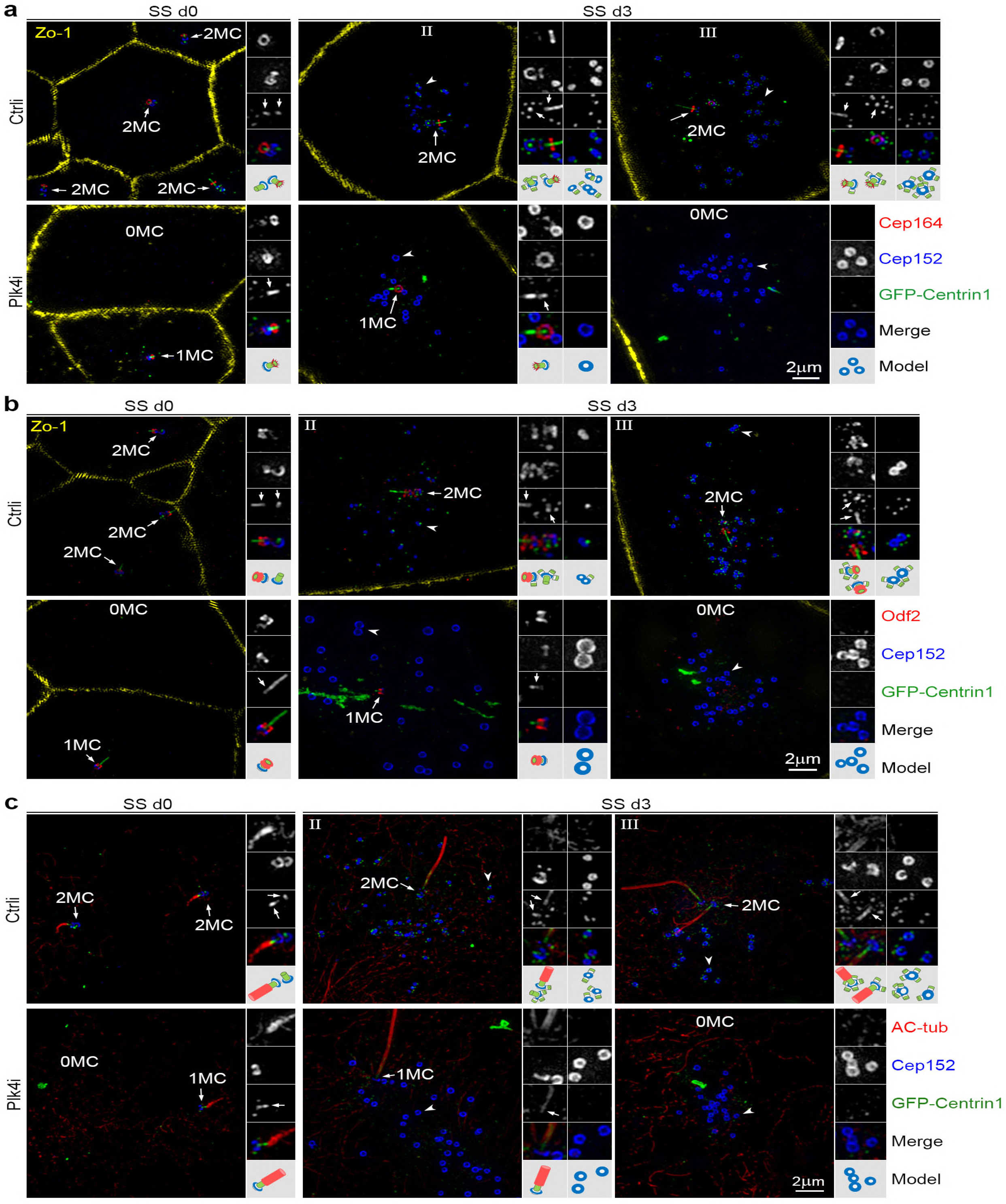
Status of the remaining MC in the Plk4-depleted mEPCs. (a,b) The remaining MC was positive for centriolar appendage proteins Cep164 and Odf2. Cultured mEPCs treated as described in Fig. 3b were immunostained to visualize Zo-1, Cep152, and Cep164 or Odf2. GFP-Centrin1, whose green fluorescence was enhanced using anti-GFP antibody and Alexa Fluor-488-conjugated secondary antibody, served as both infection and centriole markers. MCs and representative deuterosomes, denoted respectively by arrows and arrowheads, were magnified 2-fold to show details. An illustration is provided for each set of the magnified images. **(c)** The remaining MC was a basal body. mEPCs treated as described in Fig. 3b were immunostained to visualize Cep152, GFP, and acetylated tubulin (AC-tub; cilia marker). Note that the bottom region of the cilia is Centrin-positive, which explains why the Centrin-staining of the old MC frequently extends far over the centriolar appendages (Fig. 4a,b). Ciliogenesis of the young MC in stage-III control cells also explains why the MC becomes Cep164- and Odf2-positive in Fig. 4a,b.

Immunostaining using the ciliary marker acetylated tubulin (Piperno & Fuller, 1985) confirmed that a cilium already existed in the SS d0-cells (Fig. 4c) as reported (Delgehyr et al., 2015). The cilium became elongated at SS d3 (Fig. 4c). We found that the bottom region of these cilia was usually Centrin-positive (Fig. 4c). This explains the frequent observation of a Centrin-positive stick over the appendages of the old MC (Fig. 2-4), though its physiological significance remains to be clarified. In addition, we noticed that, in a portion of Ctrli-expressing mEPCs in stage III, the young MC became positive for Cep164 and Odf2 as well (Fig. 4a,b), suggesting its maturation into a basal body. Immunostaining for acetylated tubulin confirmed ciliogenesis of the young MC (Fig. 4c). This is interesting because in cycling cells the young MC needs to go through mitosis to mature into the old MC, which possesses appendages and can serve as a basal body (Loncarek & Bettencourt-Dias, 2018, Nigg & Holland, 2018). Possibly this unexpected MC maturation is rendered by the recently reported mitosis-like program in the differentiating mEPCs (Al Jord et al., 2017).

Taken together, we demonstrated that MCs are not important for deuterosome formation (Fig. 3-4). mEPCs without any MC or with only the old MC produced similar numbers of deuterosomes. Furthermore, even if the slight increase of deuterosome numbers in the control cells over the Plk4-depleted cells (4.5-4.6 deuterosomes per cell on average) (Fig. 3g) is attributed to the younger MC, this MC still cannot be considered as a prerequisite for deuterosome assembly as proposed (Al Jord et al., 2014). Our results thus support the idea of spontaneous deuterosome assembly (Fig. 1a) (Zhao et al., 2013).

Our results support the model of parallel centriole biogenesis by both MCs and deuterosomes (Fig. 1a) (Zhao et al., 2013) but not the one in which MCs are proposed to be the sole sites of procentriole nucleation (Fig. 1b) (Al Jord et al., 2014). Centriole amplifications in mTECs and mEPCs share several common features. Firstly, they both experience similar centriole amplification process, which can be classified into six stages (Fig. 1-2) (Yan et al., 2016, Zhao et al., 2013). Those in stage VI are undergoing multicilia formation (Fig. 2c) (Zheng et al., 2018). Secondly, discrete deuterosomes initially emerge as smaller ones free of procentrioles, followed by size enlargement and increased procentriole nucleation (Fig. 1-2). The model proposed by Al Jord and colleagues (Fig. 1b) (Al Jord et al., 2014) is mainly based on live cell imaging using Centrin2-GFP as marker, which makes detection of the procentriole-free deuterosomes impossible. Moreover, the spinning disk microscopy (Al Jord et al., 2014) could have missed the procentriole-associated deuterosomes in stage II due to poor optical resolution. Thirdly, both the young and old MCs start to generate procentrioles from stage II, slightly preceding the appearance of deuterosome-associated procentrioles.

Our results, however, do not necessarily contradict with the observation that the young MC can serve as a deuterosome nucleation site (Al Jord et al., 2014). In both mTECs and mEPCs Deup1 immunofluorescent signals co-localizing with those of Cep152 can sometimes be detected at both MCs in our hands (Fig. 1c) (Yan et al., 2016, Zhao et al., 2013). The MC-localized Deup1 increases in the absence of Cep63 to compensate for the function of Cep63 in the MC-dependent centriole amplification (Zhao et al., 2013). It is thus possible that some deuterosomes are nucleated from the MC-localized Deup1, possibly from both MCs, though the young MC could be more active. Hopefully future super-resolution microscopy will enable clear visualization of the deuterosome and procentriole biogenesis in living cells to allow a more precise understanding of these processes.

## Materials and methods

### Cell culture, lentivirus production, and infection

mTECs were isolated from 4-week C57BL/6J mice and cultured as described previously (Zhao et al., 2013). mEPCs were cultured as described (Delgehyr et al., 2015, Spassky et al., 2005) with some modifications. Briefly, the telencephalon was dissected from P0 mice, followed by careful removal of the meninx, choroid plexus, hippocampus, and olfactory bulb in dissection buffer (161 mM NaCl, 5 mM KCl, 1 mM MgSO4, 3.7 mM CaCl2, 5 mM HEPES, and 5.5 mM Glucose, pH 7.4) on ice. The remaining tissues were cut into small pieces and digested in freshly prepared digestion buffer (10 U/ml Papain, 0.2 mg/ml L-Cysteine, 0.5 mM EDTA, 1 mM CaCl2, 1.5 mM NaOH, and 5 U/ml DNase I in the dissection buffer) for 30 min at 37 ° C. The digestion was then stopped by adding 10% FBS. After gentle pipetting with a P1000 tip, the cells were collected by centrifugation at 1,400 rpm for 5 min at room temperature. The pelleted cells were re-suspended in the culture medium and seeded in a laminin-coated flask. Neurons were shaken off after a one-day culture and the remaining cells were further cultured to reach confluency. The cells were then transferred into the wells of laminin-coated 29-mm glass-bottom dishes (Cellvis, D29-14-1.5-N) at a density of 2×105 cells per well and were maintained in serum-free medium to induce multiciliate mEPCs.

Lentiviral productions for the RNAi experiments were performed as described previously (Zhao et al., 2013). Eighteen 10-cm dishes of HEK293T cells transfected for 48 h were used to produce the lentiviral particles, which were further concentrated to 1 ml. Radial glia-enriched brain cells isolated from three P0 mice were re-suspended into 10 ml of the culture medium (Delgehyr et al., 2015, Spassky et al., 2005) containing 60 μl of the concentrated lentiviral particles and seeded into a 75-cm^2^ flask (SS d-7). To suppress the p53-dependent apoptosis associated with centriole loss (Bazzi & Anderson, 2014), 10 μM of the p53 inhibitor, Pifithrin-α ( S2929, Selleckchem), were always included in the culture medium to sustain cell viability (Djuzenova, Fiedler et al., 2015). After 24 h of culture, neurons were shaken off and fresh culture medium was added (SS d-6). After additional six days (SS d0), the cells were serum starved to induce differentiation and assayed at SS d3.

Experiments involving mouse tissues were performed in accordance with protocols approved by the Institutional Animal Care and Use Committee of Institute of Biochemistry and Cell Biology.

### Antibodies

Secondary antibodies used for immunofluorescence (IF) were: donkey anti-rabbit conjugated with Cy3 or Dylight 405, anti-chicken conjugated with Alexa Fluor-488 or DyLight-405, anti-mouse conjugated with Cy3 or DyLight-405, anti-rat conjugated with Alexa Fluor-488, anti-guinea pig conjugated with Cy3 (Jackson ImmunoResearch), and anti-mouse conjugated with Alexa Fluor-647 (ThermoFisher Scientific). The DyLight 405-conjugated antibodies were used at 1: 200 and the remaining antibodies were at 1: 1000. Secondary antibodies used for Western blotting (WB) were HRP-conjugated goat anti-mouse and anti-rabbit antibodies (ThermoFisher; 1: 5000).

Commercial primary antibodies used were: mouse anti-Sas6 (sc-81431, Santa Cruz; IF 1:50), mouse anti-Centrin (04-1624(20H5), Millipore; IF 1:200), mouse anti-α-Tubulin (T5168, Sigma-Aldrich; WB 1:5000), mouse anti-acetylated tubulin (T7451, Sigma-Aldrich; IF 1:1000), mouse anti-Zo-1 (33-9100, ThermoFisher; IF 1:1000), rabbit anti-Centrin 1 (12794-1-AP, Proteintech; IF 1:200), rat anti-GFP (sc-101536, Santa Cruz; IF 1:50), rabbit anti-GFP (A-11122, ThermoFisher; WB 1:3000), and rabbit anti-Cep63 (06-1292, Millipore; IF 1: 200).

Rabbit anti-Deup1 (IF 1: 200; WB 1: 4000), chicken anti-Cep152 (IF 1: 300), and rabbit anti-Plk4 (IF 1: 200; WB 1: 2000) antibodies were homemade (Zhao et al., 2013). To generate antibodies against murine Cep164 or Odf2, the cDNA fragments of mouse Cep164 (NM_001081373, 1-400 aa) and mouse Odf2 (NM_001177659, 401-826 aa) were PCR-amplified from the total cDNAs of mTECs and subcloned into pET32a to express His-tagged fusion proteins. The proteins were purified by using Ni-NTA beads (Qiagen) and used as antigens. Rabbit anti-Cep164 (IF 1: 200) and guinea pig anti-Odf2 (IF 1: 200) antibodies were generated through contracted services (ABclonal) and affinity-purified.

### Immunofluorescent microscopy

Immunostaining and immunofluorescent microscopy were performed as described (Zhao et al., 2013). Briefly, mEPCs grown on glass-bottomed dishes and mTECs on Transwells were pre-extracted with 0.5% Triton X-100 in PBS for 40 sec or for 3 min, respectively, followed by fixation with 4% fresh paraformaldehyde in PBS for 15 min at room temperature. After fixation, the cells were permeabilized with 0.5% Triton X-100 in PBS for 15 min and blocked with blocking buffer (4% BSA in TBST) for 1 h at room temperature. Primary and secondary antibodies were diluted into the blocking buffer and applied to cells at room temperature for 2 h and 1 h, respectively, interspaced with 3 rounds of washing. The samples were imaged with a structured illumination microscope (GE OMX V3) with a 100×/1.40 NA oil-immersion objective lens (Olympus). Serial Z-stack sectioning was performed at 125-nm intervals. Raw images were processed for maximum intensity projection with SoftWoRx software.

### Quantification and statistical analysis

Deuterosome diameters were measured using the ‘count /size’ function of Image-Pro Plus 6.0 software (Media Cybernetics). Cells in different categories were scored manually using available original 3D-SIM images (1024×1024 pixels), each of which contained multiple cells, from three independent experiments. Quantification results are presented as mean ± SD unless otherwise stated. Differences are considered significant when *P* was <0.05 in an unpaired Student’s *t*-test using Graphpad Prism software (Graphpad Software).

## Acknowledgements

The authors thank Dr Nathalie Spassky (CNRS, France) for mEPC culture protocol and the Centre for Biological Imaging, Institute of Biophysics, CAS, for supports on 3D-SIM imaging. This work was supported by National Natural Science Foundation of China (31330045 to X.Z. and 31501092 to H.Z.), National Key R&D Program of China (2017YFA0503500), and Chinese Academy of Sciences (XDB19000000).

## Author contributions

X.Z. and X.Y. conceived and directed the project; H.Z. and Q.C. performed major experiments; Q.H. generated the homemade antibodies; X.Z., X.Y., and H.Z. designed experiments, interpreted data, and wrote the paper.

